# Dynamic marine viral infections and major contribution to photosynthetic processes shown by regional and seasonal picoplankton metatranscriptomes

**DOI:** 10.1101/176644

**Authors:** Ella T. Sieradzki, Ignacio-Espinoza J. Cesar, David M. Needham, Erin B. Fichot, Jed A. Fuhrman

## Abstract

Viruses are an important top-down control on microbial communities, yet their direct study in natural environments has been hindered by culture limitations^1-3^. The advance of sequencing and bioinformatics over the last decade enabled the cultivation independent study of viruses. Many studies focus on assembling new viral genomes^4-6^ and studying viral diversity using marker genes amplified from free viruses^7,8^. We used cellular metatranscriptomics to study community-wide viral infections at three coastal California sites throughout a year. Generation of and recruitment to viral contigs (> 5kbp, N=66) allowed tracking of infection dynamics over time and space. Here we show that while these assemblies represent viral populations, they are likely biased towards clonal or low diversity assemblages. Furthermore, we demonstrate that published T4-like cyanophages (N=50) and pelagiphages (N=4), having genomic continuity between close relatives, are better tracked using marker genes. Additionally, we demonstrate determination of potential hosts by matching infection dynamics with microbial community composition. Finally, we quantify the relative contribution of various cyanobacteria and viruses to photosystem-II *psbA* expression in our study sites. We show sometimes >50% of all cyanobacterial+viral *psbA* expression we observed is of viral origin, which highlights the proportion of infected cells and makes viruses a remarkable contributor to photosynthesis and oxygen production.

---

We sampled surface seawater in different seasons over three sites across the San Pedro Channel, California, USA: The Port of Los Angeles (POLA), Santa Catalina Island Two Harbors (CAT) and the San Pedro Ocean Time-series (SPOT). These sites represent a gradient of human impact with POLA being the most impacted and SPOT resembling open ocean conditions. In all of these sites free virus-like particles outnumber bacteria and archaea roughly 10:1 (sup. fig. 1). We examined only the 0.2-1 μm size-fraction, which includes most bacteria, archaea and some picoeukaryotes. Via assembly of metatranscriptomes, we obtained 1455 contigs longer than 5 kb of which 57 (3.9%) were characterized as viral using virSorter and virFinder (see methods). Additionally, a cross-assembly of the metatranscriptomic viral contigs with metagenomes of the same samples (N=12) yielded 9 more contigs (mean length 26,563 bp) characterized as viral. Most of the contigs represent dsDNA viruses (N= 65) as apparent from their presence in metagenomes, but one appears to be an RNA virus possibly infecting a eukaryotic host. This contig contained an RNA-dependent-RNA-polymerase whose nearest match in NCBI non-redundant database was marine Antarctic phytoplankton RNA virus PAL_E4^9^. These 66 viral contigs revealed varied patterns of presence (in metagenomes) and activity (in metatranscriptomes) in the three sites over a year (fig. 1).

**Figure 1:**
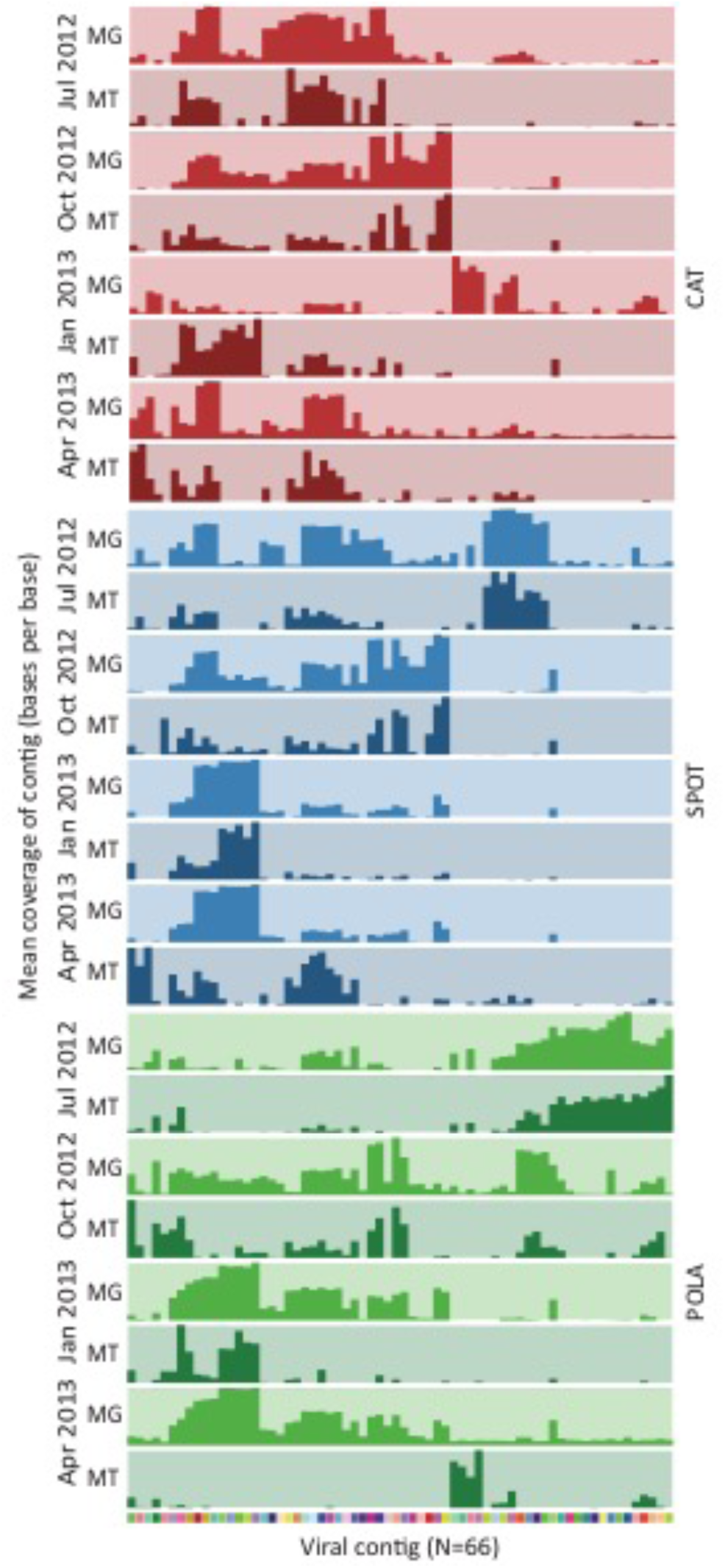
Mean coverage of 66 viral contigs across three sites (Port of LA – POLA, San Pedro Ocean Time-series – SPOT and Two harbors – CAT) and four dates (July 2012, October 2012, January 2013 and April 2013) in metagenomes (MG) and metatranscriptomes (MT). The bar heights are normalized to the highest mean coverage within the sample. Each cell in the color bar on the bottom represents a contig and corresponds with the column above it in all samples. Mean coverage was calculated excluding contig positions in the 4^th^ quartile of coverage depth which can be biased by recruitment localized to a small portion of the contig (sup. fig. 3).

Active non-synchronized viral infection would manifest as recruitment to an entire contig in both metagenome and metatranscriptomes of the same sample. We found that patterns of mean coverage from metagenomes and metatranscriptomes of our assembled viral contigs usually differed, not just between metagenomes and metatranscriptomes but also between dates and locations, implying widespread boom-bust dynamics of infection. While some variation may be due to synchronization known for some photosynthetic and heterotrophic bacteria in the ocean^10,11^ and for some of their phages^12^, this explanation is less likely as samples were collected from all sites within the same 4 hours morning-time window.

Some regional patterns were evident, e.g. some viral contigs were unique to the Port of LA (fig. 1), and that site always clustered separately from SPOT and CAT by Bray-Curtis similarity of expression of viral contigs (sup. fig. 1B). This pattern corresponds to the difference in biotic parameters between the port and the other sites (sup. fig. 2), though the port did not cluster separately in microbial community composition by 16S-rRNA (sup. fig. 1A). The latter may reflect offshore microbes brought in with the tide but less active than port organisms. Clustering by metagenomic recruitment to viral contigs did not reveal consistent patterns by site or date (sup. fig. 1C).

**Figure 2:**
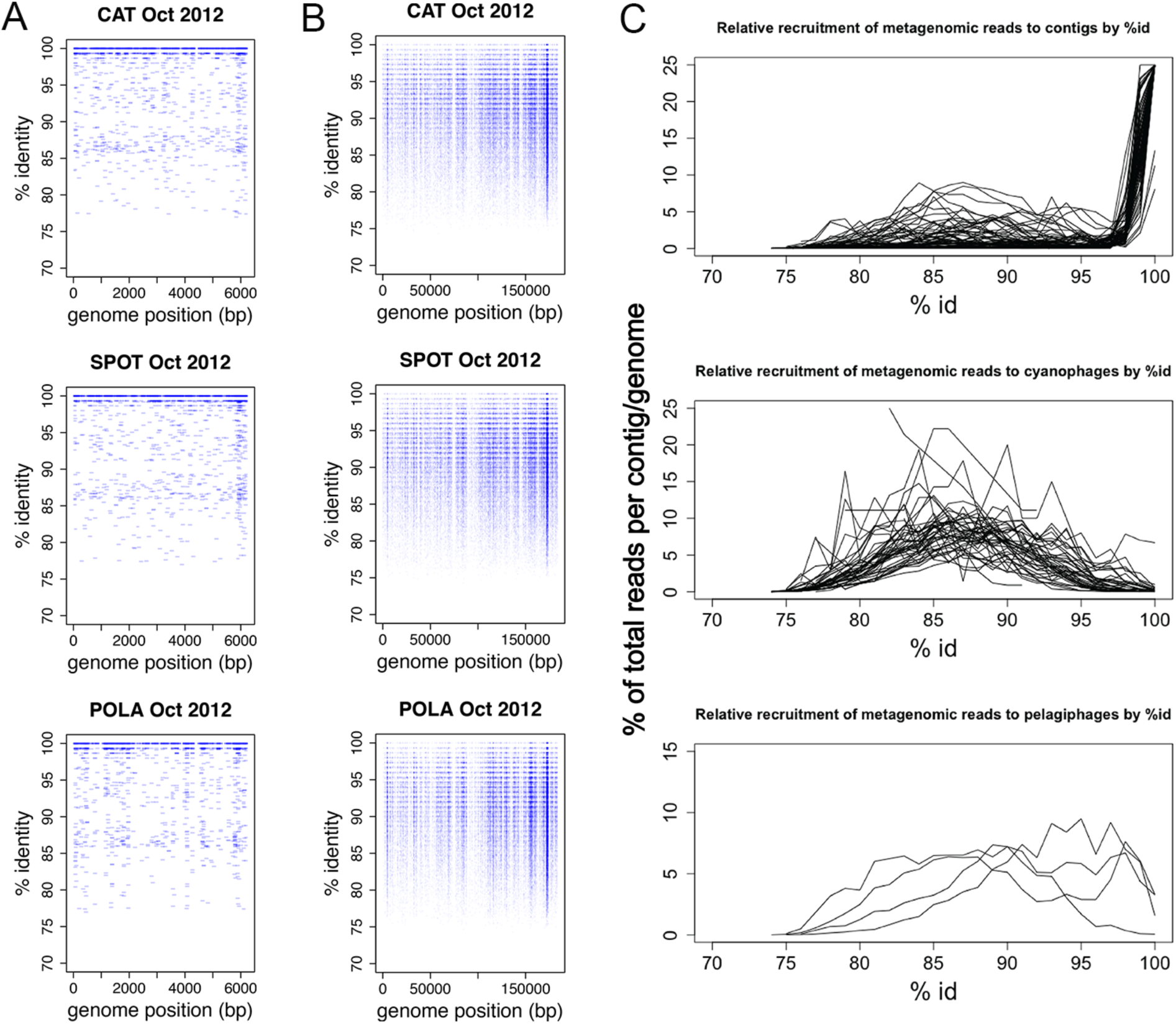
Metagenomic read recruitment to (**A**) an assembled cyanophage contig and (**B**) *Prochlorococcus* phage P-HM2 genome. Most recruitment to the assembled contig is at 99-100% identity (high density near 100% is not fully evident from the graph due to overlaps, see C), whereas P-HM2 reveals a genomic continuum. (**C**) Recruitment as a function of percent ID of reads demonstrates that assembled contigs mostly recruit at 100% ID and have few moderately close relatives (top) whereas published genomes of cyanophages reveal clouds of moderately close relatives but few matches near 100% (middle), and pelagiphages range from 100% down (bottom).

Ephemeral infections dominated the assembled landscape, as 56 out of 66 of the contigs only appeared in few metatranscriptomes, presumably reflecting sporadic infections. Persistent infections (mean coverage >= 0.75x in at least 3 out of 4 samples per site, 10 out of 66) were limited to CAT and SPOT except for one that was persistent in all three sites. Moniruzzaman et al.^13^ also recently demonstrated dominance of ephemeral dynamics in infections of marine single-cell eukaryotes during an algal bloom. Bray-Curtis dissimilarity of the viral contigs within each site was 80-100%, whereas the dissimilarity of microbial communities within site was distributed around 50-70%. High dissimilarity indicates that even within site different viruses are actively infecting in different seasons (sup. fig. 1D+E).

Moreover, assembled viral contigs appeared to be biased towards low-microdiversity (i.e. more clonal) viruses. High diversity, extremely common in marine microorganisms^14^, tends to break assemblies created with either read-overlaps or DeBruijn graphs^15,16^. We expect that low virus diversity could result from boom-bust lifestyle due to bottlenecks during “bursts”. This might lead to a method bias towards ephemerally infecting viruses. Indeed, all the viral contigs we assembled in this study appear to have many nearly identical relatives but few moderately close ones as shown by recruitment plots (most recruitment at 98-100% identity and little recruitment at 90-97%, fig. 2C), while some of the published pelagiphages had recruitment along most of the genome and high mean coverage at up to 100% identity and yet did not assemble (fig. 2C, sup. table 1).

The recruitment plots also reveal a common pattern of recruitment to short fragments near 100% identity whereas the rest of the genome or contig is only recruited to at lower percentage if at all (sup. fig. 3). This pattern highlights two issues: (1) some genes are so conserved or so often laterally transferred that their partial sequences cannot be used to identify which phage is present and (2) that mean coverage of contigs could be highly biased by these conserved regions which needs to be considered when evaluating abundance of the contigs and for coverage-based binning of genomes.

A previous report indicated that Synechococcus phage genomes occur in discrete “clouds” with a discontinuity in recruitment below ∽95% identity^17^. While this pattern exists for some cyanophage genomes, and we often saw some gaps in coverage at ∽90-95% consistent with that idea (sup. fig. 3), it is by no means the rule in our data, especially for pelagiphages (fig. 2C). We also note that widely used recruitment algorithms only map reads with a local or end-to-end match at a very high percent identity, and would therefore miss much genetic diversity that may be relevant (fig. 2B).

We were surprised not to find multiple cyanophage (especially myovirus) contigs, because such cyanophages belong to the family *Myoviridae*, some of the most common dsDNA viruses in the ocean^18^ and we know this region has a diverse community of myoviruses and cyanobacteria^7,14^. Few of the assembled viral contigs contained myoviral marker genes (e.g. capsid protein gp23) (sup. Table 2). The only assembled contig that is with high certainty from a cyanophage is a putative podovirus (see below). Recruitment of reads to published cyanophage genomes revealed the likely reason for so few such contigs: high genomic diversity (fig. 2B) which probably broke assemblies of T4-like cyanophages. We lacked assemblies despite persistent myovirus activity. We assigned translated reads identified by a Gp23-HMM (Hidden Markov Model) to published and assembled Gp23 proteins. Most versions of this marker gene from published genomes as well as the nine assembled Gp23 ORFs were expressed persistently throughout all sites and dates (sup. fig. 4). While the exact published genomes themselves were not present in our samples (fig. 2B), we posit that other T4-like cyanophages closely-related to those published are present and persistently infecting their hosts.

Matching viral contigs and hosts is challenging, but we were able to use physiological information and distributions among samples to make a likely match. Many cyanophages contain a variety of genes that maintain photosynthetic activity in the host during infection, from “spare parts” for photosynthetic reaction centers through regulation and optimization of those apparati^19^. In particular, viruses were shown to maintain photosystem II function during infection in order to supply energy to the host, as transcription of host genes is shut down during infection and PS-II proteins have a short lifetime^20,21^. Our assembled cyanophage contig contained genes coding for photosystem-II protein D1 (*psbA*) and high-light induced protein (*hli*) reportedly widespread in cyanophages^8^. The putative cyanophage from which this contig was derived was actively transcribed (presumably infecting its host) in all three sites only in October 2012 (fig. 4A). The cyanobacterial community by 16S-rRNA was dominated in October by two operational taxonomic units (OTUs): one *Synechococcus* and one *Prochlorococcus*. Both OTUs were present at SPOT and CAT in October, but only *Synechococcus* was also present at POLA (fig. 4B).

Thus, we propose that this assembled contig is from a phage that infects *Synechococcus* OTU 10 which has a 16S sequence over the amplified region 100% identical to *Synechococcus* CC9902 of clade IV. On a phylogenetic tree of PS-II D1, translated PS-II D1 of this phage clustered closely with a different phage isolated on *Synechococccus* (sup. fig. 5).

**Figure 3:**
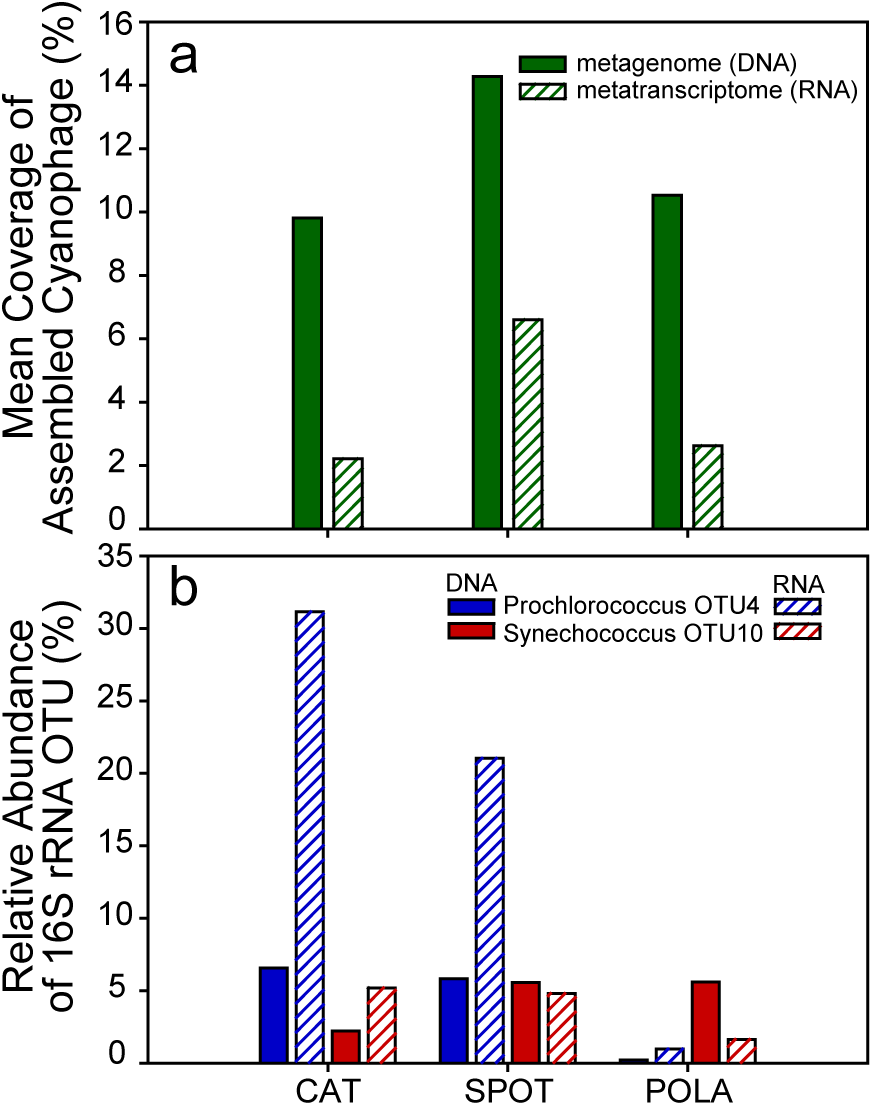
Presence and activity of the assembled cyanophage and its potential hosts in October 2012: (**A**) Mean coverage (quartiles 1-3) of assembled cyanophage (**B**) OTU relative abundance by 16S-rRNA of the two most abundant cyanobacteria OTUs in order: *Prochlorococcus* in DNA, *Prochlorococcus* in RNA, *Synechococcus* in DNA, *Synechococcus* in RNA. Note the near-absence of *Prochlorococcus* in POLA, in contrast to *Synechococcus* and the phage, leading us to infer the phage infects *Synechococcus*.

Because viruses and hosts both code for photosynthetic functions, a comparison of viral and host-coded contributions to activity is possible. Sharon et al.^22^ previously showed viral *psbA* gene can outnumber cyanobacterial *psbA* genes in metagenomes from the Mediterranean, and showed viral gene expression is evident. We extended this to quantitatively partition gene expression into bacterial contribution from *Synechococcus* and *Prochlorococcus* and viral contribution from cyanomyoviruses and cyanopodoviruses, as evident from HMM-placed translated reads onto our PS-II D1 phylogenetic tree. We found *psbA* transcripts of T4-like cyanomyovirus origin generally accounted for roughly 50% of cyanobacterial and cyanophage *psbA* transcripts. *Prochlorococcus* transcripts were almost always comparable to the T4-like contribution. On several occasions, the viral version exceeded the cyanobacterial version in read count (fig. 4).

**Figure 4:**
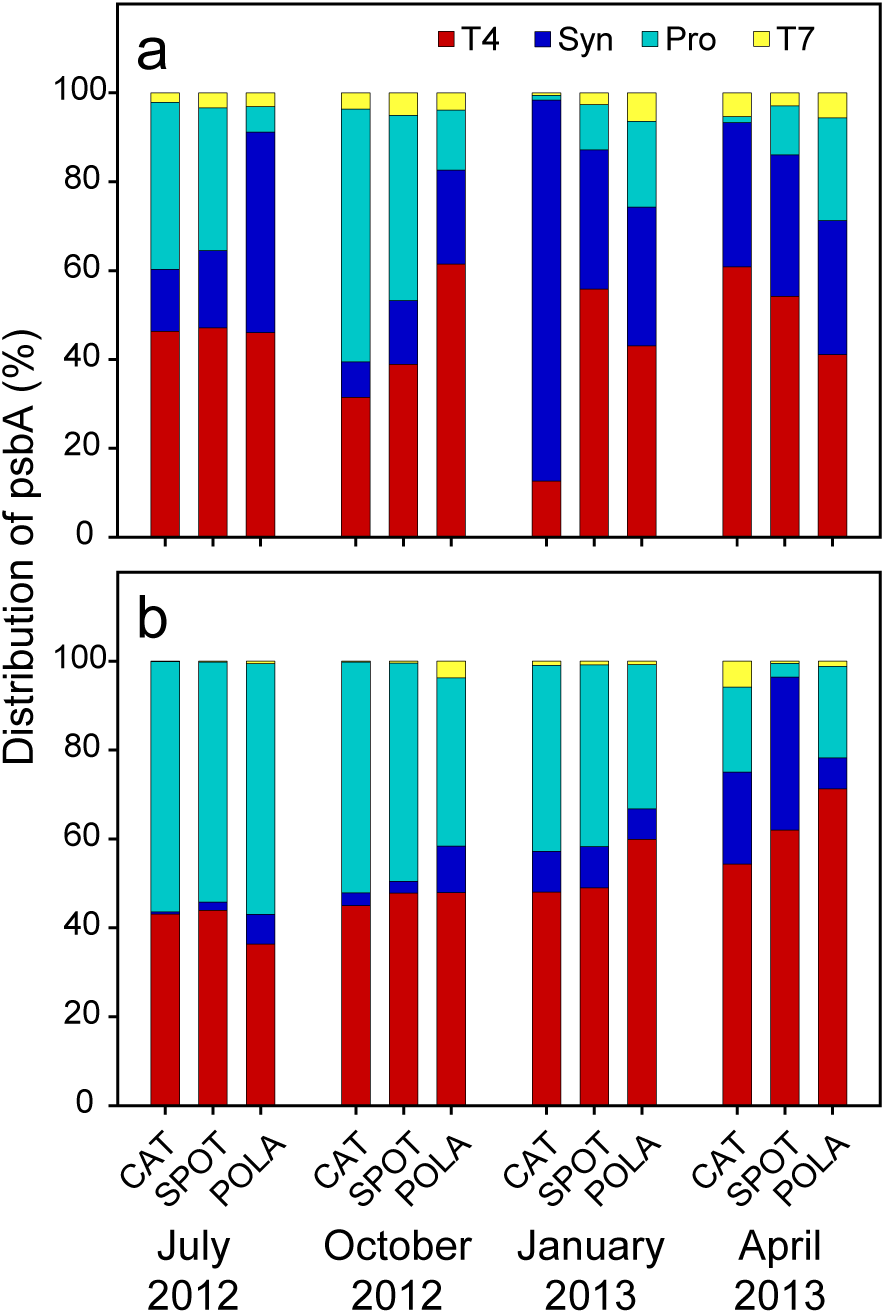
Distribution of psbA of T4-like phages, *Synechococcus*, *Prochlororoccus*, and T7-like phages in (A) metagenomes and (B) metatranscriptomes.

We can roughly estimate the proportion of infected cyanobacteria from our *psbA* data and compare it to previously published estimates. For cyanobacteria in marine systems, the highest estimates of infection are roughly 50-60% infected at any given time^2,17,23,24^. One consideration when calculating the proportion of infected cyanobacteria is that during host infection, the number of phage mRNA of *psbA* increases quickly during early infection until it becomes the exclusive source of *psbA* transcripts in the cell^20,21^. Another consideration is that, regardless their source, host or virus, the abundance of *psbA* transcripts is comparable in infected and uninfected cells^23^. What we observe in the sample is a comparable contribution of T4-like phages and cyanobacteria (fig. 5 D) at a ratio of 1.2±0.6 (mean ± standard deviation) phage/cyanobacteria, which suggests that on average about half of the cyanobacteria are infected. This is in accordance with the high end of published estimates, confirming that infection is an important part of cyanobacterial ecology.

In both metagenomes and metatranscriptomes, there is minor consistent recruitment to T7-like cyanopodovirus psbA. However, in every sample the contribution of T7-like cyanopodoviruses was very low compared to that of T4-like cyanomyoviruses. This could be due to the more specific host range reported for cyanopodoviruses compared to cyanomyoviruses^25-27^. As T4-like and T7-like cyanophages are reported to be strictly lytic^28^, their presence in metagenomes results from late infection genomic copies or virions within host cells, pseudolysogeny or phages that adsorbed to cells or particles.

Extending metatranscriptomics methods as recently applied to marine eukaryotic viral infection^13,29,30^, we show the power of multiple approaches to track viral infection and dynamics within the broad picoplankton community, using metatranscriptomes of the cellular fraction, with particular examples in the cyanobacteria. Use of marker genes is especially important to study viruses with many close relatives in the same environment (whose contigs assemble poorly), whereas assemblies are useful for tracking ephemeral, more clonal viruses. The observed infection dynamics can sometimes be used in combination with microbial community structure and viral marker genes found within contigs to deduce a host. Use of metagenomes and metatranscriptomes provides an insight into quantifiable viral contribution to photosynthesis and to estimating the fraction of infected cyanobacteria.

## Methods

### Sample collection

Surface seawater was collected by bucket on 7/15/2012, 10/19/2012, 1/9/2013 and 4/24/2013 in three locations: The Port of Los Angeles (33°42.75’N 118°15.55’W), the San Pedro Ocean Time-series (33°33.00’N 118°24.01’W) and Two Harbors, Santa Catalina Island (33°27.18’N 118°28.51’W). Duplicate samples of 20 liters were filtered in each location through an 80 μm mesh followed by a glass fiber syringe prefilter (Gelman, 4523) which collected the >1 μm size fraction and a 0.2 μm PES Sterivex filter (Millipore, SVGPB1010) which collected the free-living size fraction. RNAlater (Thermo-Fisher, AM7020) was added to each filter and filters were flash frozen no more than 5 minutes post-filtration.

### Library preparation

DNA and RNA were extracted simultaneously from Sterivex filters by bead-beating followed by an AllPrep kit (Qiagen, 80204). An internal standard (ERCC RNA Spike-In Mix, Thermo-Fisher 4456740) was added into the lysate after bead-beating for quality assurance. RNA was enriched for mRNA with RiboZero (Illumina, MRZB12424). Resulting mRNA was reverse transcribed using SuperScript-III (Invitrogen, 18080-051). DNA and cDNA were sheared with Covaris m2 and size-selected for products larger than 300 bp. RNA libraries were prepared and barcoded using NEBNext Ultra Directional RNA library Prep Kit for Illumina (E74205). DNA libraries were prepared and barcoded with Ovation UltraLow Library Prep V2 (Nugen, 0344).

Metagenomes were sequenced on Illumina HiSeq 2x125 bp or 2x150 bp. Metatranscriptomes were sequenced on Illumina HiSeq 2x250 bp.

### Read processing and assembly

Raw metagenomics and metatranscriptomics reads were quality trimmed and filtered with Trimmomatic version 0.33 with parameters LEADING:20 TRAILING:20 SLIDINGWINDOW:15:25^31^. Metatranscriptomic reads were merged with PEAR^32^, using the default settings and residual ribosomal reads as well as the internal standard were removed informatically. Merged reads from each sample separately were assembled with Megahit.

Contigs smaller than 2kbp from all samples were co-assembled with Newbler^33^ version 2.9 (Roche) (minimum overlap 40bp minimum id 99%) and contigs larger than 2kbp from all samples were co-assembled with minimus2^34^ (minimum overlap 40bp minimum id 99%). Only contigs larger than 5 Kbp were further analyzed.

### Identification and annotation of viral contigs

Viral contigs were identified by VirSorter^35^ using RefSeq on the CyVerse platform and only contigs classified as category 1 or category 2 were considered. In addition, the contigs were ranked using VirFinder^36^ (rank >=0.95). Prodigal^37^ was used to predict ORFs in those contigs, and the amino acid sequences were searched against the nr database (August 12^th^ 2016) using blastp^38^ and a maximum E-value 10^−5^. The annotations were used to verify viral contigs from the VirFinder results. Contigs were verified to be non-chimeric by even recruitment.

Quality filtered metagenomic and metatranscriptomic reads were mapped back to these contigs with Bowtie2 version 2.2.6 using the default settings and the expression patterns were identified and visualized with Anvi’o^39^ version 2.1.0.

### Microbial community composition analysis

The V4-V5 regions of the 16S-rRNA coding gene were amplified from DNA and cDNA from all samples using the 515-N-F and 926-R primers, and sequenced on an Illumina MiSeq 2x300 bp (UC Davis genome center) along with a mock community as described in Parada et al.^40^.

The ends of resulting reads were trimmed with PRINSEQ^41^ to a quality score higher than 20. The trimmed reads were merged with USEARCH7^42^ allowing for 3 mismatches in the overlap region. Retained assembled reads were clustered with mothur^43^ version 1.38.0 according to the MiSeq and classified with SILVA version 119. Bray-Curtis dissimilarity and dendrograms were calculated and plotted with R package vegan^44^.

### Analysis of PS-II D1 protein sequences

A curated set of PS-II D1 amino acid sequences of myoviruses, podoviruses, cyanobacteria and eukaryotes (chloroplast) from Pfam^45^ and RefSeq release 80 was downloaded. All sequences of marine viral PS-II D1 were retained in addition to sequences of bacterial and eukaryotic taxa that were identified in the 16S-rRNA community composition. One of the assembled contigs contained a psbA gene coding for PS-II D1. The translated amino acid sequences were added to the set of proteins.

Merged reads from the metatranscriptomes and unmerged forward reads from the metagenomes were aligned with blastx^38^ against this set demanding an e-value of 10^−5^. The reads that passed the filter were translated using bioPython^46^ into amino acids according to the reading frame indicated by the blastx start and end values.

Following the protocol used in Ignacio-Espinoza et al.^47^ total of 158 sequences were aligned with mafft^48^ version 7.305b with parameters set to globalpair, gap open penalty 1.5, gap extension penalty 0.5 and scoring matrix BLOSUM30. Informative blocks were identified using Gblocks^49^ version 0.91b with a minimum block length 5, blocks represent at least half of the sequences and allowing gaps (b3=50, b4=5, b5=h). The blocks were used to build a maximum likelihood phylogenetic tree using RAxML^50^ (best of 20 trees, gamma model and WAG substitution matrix). A hidden Markov Model (HMM) of the same set was also built with hmmer 3.0^51^. The translated metagenomics and metatranscriptomics amino acid sequences were searched using the HMM and a threshold of e-value 10^−5^. A total of 190,928 translated metatranscriptomics reads and 72,292 metagenomics reads from all samples remained after this step. Those reads were locally aligned to the HMM using hmmer 3.0 function hmmalign and placed into the phylogenetic tree using pplacer^52^ version v1.1.alpha17 (sup. fig. 6).

### Analysis of gp23 protein sequences

Metatranscriptomic and metagenomics reads were searched against a set of T4-like clusters of orthologous groups (COGs) with an E-value threshold of 10^−5^. 89,768 metatranscriptomic reads and 134,995 metagenomic reads were annotated as gp23. An HMM of gp23 was built as described previously and translated reads were searched and placed with pplacer. The tree was visualized by the Interactive Tree Of Life (iTOL)^53^.

### Recruitment to phage genomes

The four currently available full pelagiphage genomes were downloaded from NCBI and concatenated with assembled viral contigs from metatranscriptomes the metagenomes as well as with published cyanophage genomes downloaded from NCBI RefSeq. Metagenomic and metatranscriptomics reads were searched against the genomes dataset with blastn default settings. For metagenomes only hits longer than 100bp were retained, and for metatranscriptomes only hits longer than 200bp. Hits were then plotted against the genomes using R^54^.

### Data availability

All data can be found on EMBL-ENA under project number PRJEB12234. Raw metatranscriptomics sequences accession numbers are ERS1864892-ERS1864903, and negative control library sequences accession number is ERR2089009. Raw metagenomic sequences accession numbers are ERS1869885-ERS1869896 and negative control accession number is ERS1872073. Assembled viral contigs accession numbers are ERZ474118-ERZ474183.

## Acknowledgements

The authors would like to thank R. Sachdeva, N. Ahlgren, A. Parada, L. Berdjeb, E. Graham, M. Lee, J. Ren, F. Sun and T. Delmont for insightful discussions and advice on bioinformatics analyses. We thank Catherine Roney-Garcia, the Sundiver crew and the USC Wrigley Institute of Environmental Studies for logistic support. This work was supported by NSF grant 1136818, Gordon and Betty Moore Foundation Marine Microbiology Initiative grant GBMF3779 and Norma and Jerol Sonosky summer fellowship to E.T.S.

